# Behavioral training of marmosets and electrophysiological recording from the cerebellum

**DOI:** 10.1101/683706

**Authors:** Ehsan Sedaghat-Nejad, David J. Herzfeld, Paul Hage, Kaveh Karbasi, Tara Palin, Xiaoqin Wang, Reza Shadmehr

## Abstract

The common marmoset (*Callithrix Jacchus*) is a promising new model for study of neurophysiological basis of behavior in primates. Like other primates, it relies on saccadic eye movements to monitor and explore its environment. Previous reports have demonstrated some success in training marmosets to produce goal-directed actions in the laboratory. However, the number of trials per session has been relatively small, thus limiting the utility of marmosets as a model for behavioral and neurophysiological studies. Here, we report the results of a series of new behavioral training and neurophysiological protocols aimed at increasing the number of trials per session while recording from the cerebellum. To improve the training efficacy, we designed a precisely calibrated food regulation regime that motivated the subjects to perform saccade tasks, resulting in about a thousand reward-driven trials on a daily basis. We then developed a multi-channel recording system that used imaging to target a desired region of the cerebellum, allowing for simultaneous isolation of multiple Purkinje cells in the vermis. In this report, we describe (1) the design and surgical implantation of a CT guided, subject specific head-post, (2) the design of a CT and MRI guided alignment tool for trajectory guidance of electrodes mounted on an absolute encoder microdrive, (3) development of a protocol for behavioral training of subjects, and (4) simultaneous recordings from pairs of Purkinje cells during a saccade task.

**New and Noteworthy:** Marmosets present the opportunity to investigate genetically based neurological disease in primates; in particular, diseases that affect social behaviors, vocal communication, and eye movements. All of these behaviors depend on the integrity of the cerebellum. Here, we present training methods that better motivate the subjects, allowing for improved performance, and also present electrophysiological techniques that precisely target the subject’s cerebellum, allowing for simultaneous isolation of multiple Purkinje cells.

*In our parks, are there any trees more elegant and luxurious than the Purkinje cell from the cerebellum?* Santiago Ramon y Cajal

## Introduction

The common marmoset (*Callithrix jacchus*) has gained attention for neurophysiological investigation in recent years because of its potential for transgenic manipulation (Sasaki et al., 2009;Kishi et al., 2014;Miller et al., 2016). Marmosets are small, New World primates with no known lethal zoonotic diseases that are transmittable to humans (Wakabayashi et al., 2018). Behaviorally, they share important attributes with us: they are social and live in family units, they are vocal and rely on species-specific sound production for their communication, and they are visual and use saccadic eye movements to explore their environment. Notably, for a primate they have a particularly short gestation period (∼ 5 months) and regularly give birth to twins or triplets. Thus, they have a high breeding efficiency with potential for germline transmission of genetically modified models.

Indeed, marmosets research is benefiting from transgenic (Sasaki et al., 2009), gene-editing (Kishi et al., 2014), and optogenetic tools that target specific neurons (MacDougall et al., 2016). Thus, marmosets have the potential to become a model system for investigating cognitive and social behaviors (Takahashi et al., 2017;Mustoe et al., 2015), auditory (Wang, 2018) and visual perception (Solomon and Rosa, 2014), as well as neural control of vocalization (Roy et al., 2011;Eliades and Wang, 2013;Eliades and Miller, 2017) and eye movements (Mitchell et al., 2014;Johnston et al., 2018).

Many aspects of social behavior, vocalization, and eye movements depend on the integrity of the cerebellum. For example, children who suffer from autism spectrum disorder (ASD), a developmental disorder that leads to impairments in social and communication skills, exhibit anatomic abnormalities in their cerebellum, including reduced number of Purkinje cells (Whitney et al., 2008), particularly along the vermis (Murakami et al., 1989;Hashimoto et al., 1995;Courchesne et al., 2001;Scott et al., 2009). Notably, in children with ASD, we found that damage to the cerebellum is prevalent in lobule VI and parts of lobule VIII (Marko et al., 2015), regions that in the macaque are critical for control of saccadic eye movements (Takagi et al., 1998;Barash et al., 1999). Control of saccades in healthy subjects shows exquisite sensitivity to decision related variables such as reward prediction error (Sedaghat-Nejad et al., 2019), and history of effort expenditure (Yoon et al., 2018). Thus, marmosets may present a particularly good opportunity to investigate the role of the cerebellum in eye movements, social communication, and neurological diseases such as ASD.

However, despite these attractive features, there is concern that marmosets may be difficult to train for studies that investigate the neural basis of goal-directed behavior in a laboratory setting. This concern is due to the fact that in the current literature, the number of trials that these animals can perform within a session appears to be quite limited. For example, one recent study that trained marmosets to perform saccades found that on average, they performed 80-100 trials per session (Johnston et al., 2018), while another study reported 300-800 trials per session (Mitchell et al., 2014).

These numbers are on the lower bound of what is needed to identify task-related neurons in the cerebellum. For example, identification of an eye-related Purkinje cell requires 200-300 trials to measure complex spike tuning, and an additional 500-1000 trials to measure the simple spike response to saccadic eye movements (Soetedjo and Fuchs, 2006;Soetedjo et al., 2008). As a result, the relatively low numbers of trials per session reported earlier makes it unclear whether marmosets can serve as a model for neurophysiological studies of operant conditioned behavior.

Over the past four years, our team has been building a new marmoset laboratory, aiming to develop behavioral and neurophysiological protocols that allow for electrophysiological recording from the cerebellum of head-fixed animals. From a behavioral perspective, our goal was to ask whether the subjects could be motivated to produce a sufficiently large number of rewarded trials on a daily basis in the head-fixed configuration. From an electrophysiological perspective, our aim was to produce precise targeting of a region of the cerebellum, thus providing the means to simultaneously isolate multiple Purkinje cells using high-density electrodes.

To solve the training problem, we designed a carefully calibrated food regulation regime that motivated the subjects to perform a saccade task. To solve the electrophysiological problem, we developed a high-precision, acute multi-channel recording system that resulted in simultaneous recording from multiple Purkinje cells within the cerebellar vermis.

In this report, we provide an account of our experience, both in terms of procedures that failed and those that succeeded, going from initial behavioral training through neurophysiological recording. We begin with the design of a CT guided subject-specific titanium printed head-post, one that may be the first of its kind in the marmoset. We then describe the training and the MRI guided electrode alignment procedures. We conclude with examples of simultaneously isolated pairs of Purkinje cells, a difficult accomplishment only once before reported in the awake behaving primate (Medina and Lisberger, 2007).

## Methods

### Subjects

The procedures were carried out on three marmosets: two female (subject B, 420g, 11 years old; subject M, 350g, 4 years old) and one male (subject R, 400g, 4 years old). All three subjects were born and raised in a colony that one of the authors (XW) has maintained at the Johns Hopkins School of Medicine since 1996. The experimental procedures were evaluated and approved by Johns Hopkins University Animal Care and Use Committee in compliance with the guidelines of the United States National Institutes of Health.

### Design of head-post and recording chamber

Recording from Purkinje cells of the cerebellum imposes particular demands on stability: one needs to not only isolate a neuron, but also establish identity of that neuron via presence of both simple and complex spikes, which requires movable electrodes and longer periods of recordings. In order to stabilize the head during behavioral sessions, the current technique is to build an acrylic based thick “helmet” atop the skull, which then serves as a base that holds two or more metal posts (Lu et al., 2001a). The glue-inserted metal posts are attached to bars that are fixed to a primate chair, thereby holding the head in a fixed position. A more recent method is a halo-like device that holds the skull inside a ring (Johnston et al., 2018). Both approaches have been used for recordings from the cerebral cortex. However, reaching the cerebellum requires open access to the posterior parts of the skull, making the halo approach less desirable. Furthermore, we aimed to produce an approach that reduced disruption of the underlying skin and muscles, resulting in a design that allowed for skin coverage of the skull and integrity of the temporalis muscles. This led us to develop a titanium based head-post and chamber design that eliminated use of the acrylic based helmet, and maximized the open area in the posterior part of the skull (Fig. 1).

**Fig. 1.**
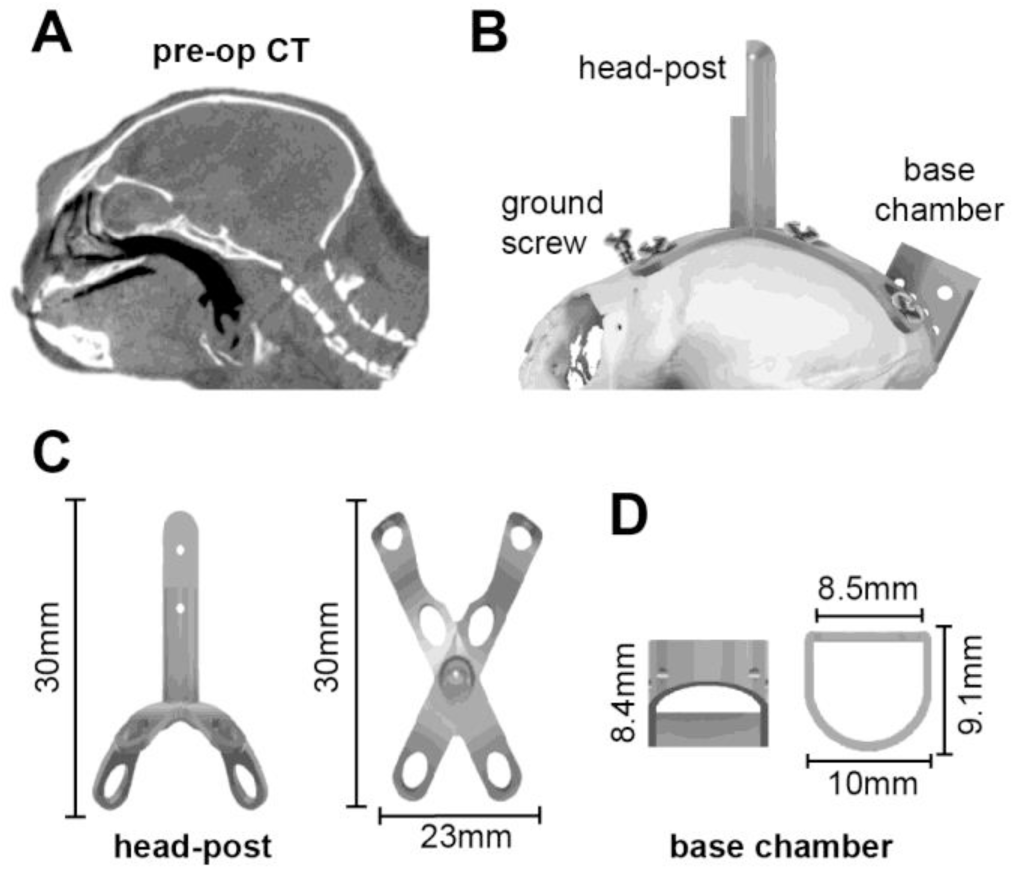
Design of the head-post and base chamber. **A**. Preoperative CT image that we used to build a geometric model of the surface of the skull. **B**. Model of the head-post and chamber fitted to the CT-based model of the animal’s skull. The screws, head-post, and chamber are titanium. **C**. The head-post (2.39 g), designed to precisely fit the curved geometry of the subject. **D**. Base chamber (0.48g).

To prepare for head-fixation, we designed a head-post based on the specific geometry of the surface of each subject’s skull and manufactured it using 3D printed titanium. We began with a preoperative CT (Fig. 1A), and used it to build a 3D model of the skull (Fig. 1B). This was done using 3D Slicer, an open-source image analysis and visualization software (Fedorov et al., 2012). The 3D skull model was then imported into a CAD environment (SolidWorks and Autodesk Fusion 360), which informed the design of an X-shaped titanium head-post (Fig. 1B and Fig. 1C), a titanium base recording chamber (Fig. 1D), and a plastic protective chamber cap. The head-post and base chamber designs were optimized to remain light weight and within the anatomical constraints of the frontal eye ridge and the lateral ridges on the skull, allowing us to minimize damage to the temporalis muscle attachment. The head-post and base chamber were 3D printed with laser melted grade 5 titanium (6AI-4V, Sculpteo.com), producing a lightweight (3 g), biocompatible, corrosion resistant structure. The protective cap that covered the chamber was 3D printed with PLA plastic filament (0.5 g).

### Surgery

Implanting the head-post and chamber was conducted by a surgical team led by lab members, and aided by veterinary technicians and veterinarians. The animal was anesthetized with Alfaxalone (12 mg/Kg), then intubated with #2.0 uncuffed endotracheal (ET) tube, and administered isoflurane (1-1.25%, adjusted as needed) to maintain the anesthetic plane, as well as atropine and dexamethasone via intramuscular (IM) injections. The subject was then head-fixed by inserting bilateral ear bars into the osseous ear canal, which were fastened to a stereotaxic frame, while a bite bar was slid into place with a forehead clamp attachment to ensure that the head was positioned appropriately and securely with respect to the chest. 2% Lidocaine and 1:100000 epinephrine was then injected into the subcutaneous space beneath the scalp, and the operative region was treated with alternating cycles of diluted chlorhexidine and alcohol and at the very end sprayed with betadine. Next, we used a surgical marker to draw an outline for a midline incision on the scalp, extending 2mm posterior to the supraorbital fat pad and extending to the occipital ridge. We then made the incision using a scalpel and the skin was separated from the underlying fascia as well as the fascia from the underlying muscle. From here, the surface of the skull was prepared by removing residual tissue using curettes and 3% hydrogen peroxide solution, while the head-post was positioned for implantation. The temporalis muscle was pushed backed from its bony insertions on the spots that intersected with the head-post.

To attach the head-post, phosphoric acid etchant was applied for 15-20 seconds and then rinsed. Next, a thin layer of Optibond Solo Plus (Kerr Corp.) was applied to the surface of the skull, covering the entire area to be occupied by the head-post, cranial ground screw, and base chamber. Optibond Solo Plus was also applied to head-post X-shaped legs and lower half of the base chamber. After application, the Optibond was cured using a UV gun. The screw locations were then marked and drilled using drill bit #54 (1.4 mm diameter), with a stopper limiting the hole depth to 1.1mm. NX3 dual-cure dental cement (Kerr Corp.) was applied to the center of the bottom of the head-post, placed on the skull, and then cured using UV light.

Titanium screws (self tapping, 2.0×4 mm, 0.04 g, or 2.0×5 mm, 0.05 g) were then used to affix the legs of the head-post onto the skull. A cranial ground screw (titanium, self tapping 2.0×8 mm, 0.08 g) was screwed into the prepared hole (Fig. 1B). We confirmed that screw depth into the skull did not exceed 1.1 mm. Next, NX3 cement was applied inside head-post screw holes, and around the ground screw to ensure structural integrity of the implanted screws.

NX3 cement was also applied to the bottom edge of the base chamber and the base chamber was placed between the rear legs of the head-post (Fig. 1B) so that it was approximately located above the visual cortex and directed straight down. NX3 cement underwent a UV light curing. A final layer of NX3 cement was applied over the whole structure and was smoothed using Parafilm tape, and then cured with UV light.

The total weight of the NX3 cement, the installed apparatus, and the screws was less than 8.5 g. With the implantation complete, the temporalis muscle and fascia were replaced to their original positions. Finally, the two skin flaps were re-attached together via sutures using a 0/4 single filament thread. As a result, the surgical procedure produced an animal in which the skin covered almost the entire skull, except for small regions dedicated to the frontal ground screw, the main bar of the head-post, and the base chamber.

At this point anesthesia was discontinued and the ET tube was removed. The animal was monitored as consciousness and motor function were regained. A second dose of Dexamethasone (0.25 mg/Kg) was administered via IM injection 12 hours after the first dose. Recovery was monitored with daily postoperative check-ups for 13 days, including inspection of the surgery site for infection, checking for integrity of the sutures, recording of vital signs, feeding, and the administering of drugs.

### Establishing a chamber-based coordinate system for the cerebellum

Because of the small size of the skull and our chamber design, we found it difficult to attach a micromanipulator to the head-post. Given that the micromanipulator was external to the head-post, we were concerned that we may not be able to accurately target and reproducibly approach a desired location in the cerebellum. This led us to design an electrode guide system that we integrated into the chamber design (Fig. 2 and Fig. 3). Our problem was that during surgery, the base chamber was placed by hand on the skull, and therefore its location with respect to the cerebellum was unknown. Furthermore, post-surgical CT or MRI imaging of the implanted animal was possible but problematic because the titanium pieces generated substantial artifacts, making it difficult to precisely localize the base chamber.

**Fig. 2.**
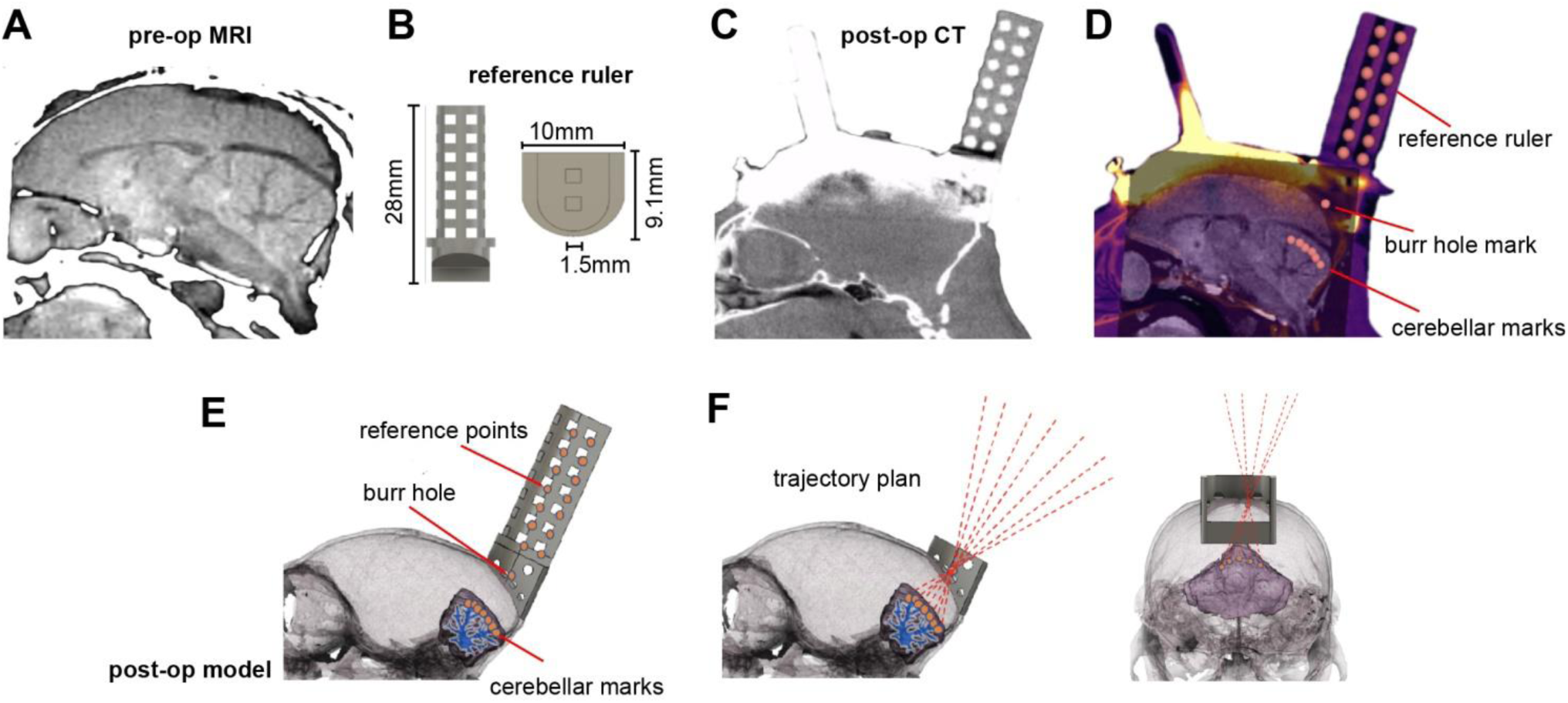
Development of a chamber-based geometric model to define electrode trajectories and specify location of the burr hole. **A**. Pre-operative MRI image used for identifying the desired regions of interest in the cerebellum. **B**. Reference axis ruler (1.104g) that was inserted into the base chamber before post-op CT imaging. **C**. Post-operative CT image after the surgical installation of the head-post, base chamber, and reference ruler. While the reference ruler is clearly visible, the titanium head-post and chamber have produced significant artifacts. **D**. Co-registered pre-operative CT, post-operative CT, and pre-operative MRI. Markers identify points on the reference axis and points in the cerebellum. A mark identifies the burr hole location. **E**. A 3D model that has co-registered the skull, cerebellum, chamber, and reference ruler. **F**. Using the 3D model, we drew trajectories that began at points of interest in the cerebellum, converged on a single 1.5mm diameter burr hole on the skull, and then diverged beyond the chamber as cylinders within the guidance tool. These trajectories represented desired electrode paths.

**Fig. 3.**
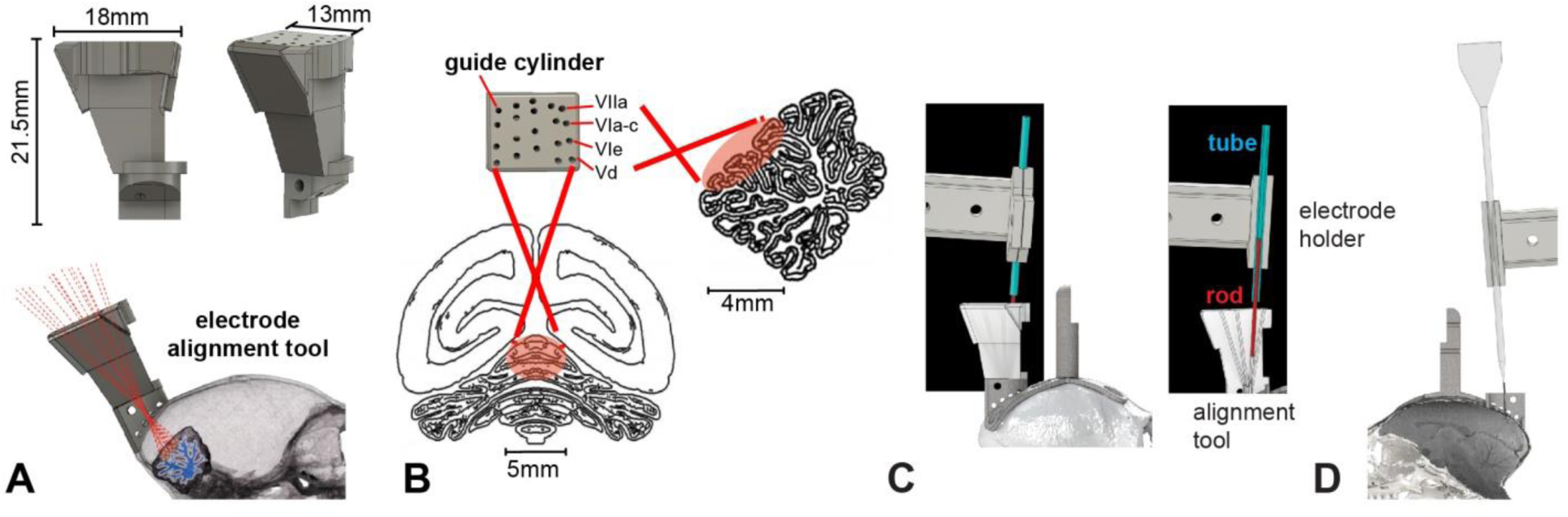
Design of the electrode trajectory guidance tool. **A**. The alignment tool was designed to allow precise placement of the electrode along each of the 20 desired trajectories that converged onto various cerebellar destinations. The tool consisted of a 3D printed block (2.03g) that had a single cylinder for each of the desired electrode trajectories. The figure shows a model of the tool with 20 guide cylinders that align with the recording regions of interest and converge through the single craniotomy burr hole. **B**. Guide cylinder trajectories with posterior and sagittal views of the recording range on the cerebellum. The labels identify the cerebellar lobule that the trajectory is aimed toward. **C**. Alignment of the electrode with the desired trajectory. We installed the alignment tool in the base chamber and then placed a rod in the cylinder corresponding to the desired trajectory. The image on the right shows a section view of the alignment tool. We then placed a tube in the electrode holder (the same holder that would later hold the electrode), and attached the electrode holder to the microdrive. The microdrive was then maneuvered by a micromanipulator so that the axis of motion of the microdrive holding the tube was aligned with the rod. **D**. An image of the electrode and the electrode holder aligned to advance along the desired trajectory.

To solve this problem, we designed a CT visible reference ruler (Fig. 2) that we inserted into the base chamber. The reference ruler consisted of 7 layers of a 1.5mm marked grid system, 3D printed with PLA plastic filament (Fig. 2B). With the reference ruler inserted into the chamber, we performed a post-operative CT scan (Fig. 2C). This produced a clear image of the ruler, but with artifacts from the titanium. We then co-registered pre-operative CT (Fig. 1A) and post-operative CT (Fig. 2C) using bony areas and co-registered pre-operative CT (Fig. 1A) and the pre-operative MRI (Fig. 2A). Fixing pre-operative CT between 2 registrations produced a co-registered image of all three images. The result was a full representation of the subject’s head, including the reference ruler, the skull, and the brain (Fig. 2D).

We used the ruler, chamber, and skull geometry to plan electrode trajectories in order to arrive at 20 distinct points of interest in the cerebellum. In this design, all 20 trajectories traveled through a single 1.5 mm diameter burr hole. To calculate each trajectory, we placed 21 points at various positions on the reference ruler (red marks, Fig. 2D), 1 point on the surface of the skull for a craniotomy burr hole (burr hole point, Fig. 2D), and 20 points at various locations along lobules V, VI, and VII of the vermis (cerebellar marks, Fig. 2D). The point representing the burr hole was placed at 1.1 mm lateral to the midline in order to avoid the superior sagittal sinus.

The various points were then imported into a CAD environment, along with a model of the brain that we generated with 3DSlicer from the MRI images (Fig. 2E). We aligned the imported points to their respective targets, guided by the co-registered image. The process relied on the alignment of all reference axis ruler points from the co-registered image (Fig. 2D) to the reference ruler within the model environment (Fig. 2E). This automatically aligned the burr hole and points on the cerebellum to the skull and brain models, creating an accurate representation of the distances and geometry of the desired cerebellar points, the desired burr hole, and base chamber.

Using the coordinates of the various points we created a set of 20 trajectories that originated from the desired recording locations in the cerebellum, converged through the single burr hole, and traveled out past the base chamber (Fig. 2F). Each line in 3D space represented a trajectory that an electrode would travel to reach a given cerebellar destination.

### Design of an electrode guidance tool

Based on the electrode trajectories that we had defined in the post-op model (Fig. 2F), we designed a tool that would be attached to the base chamber and provide alignment of the electrode to the specific trajectory that traveled through the burr hole and arrived at the desired cerebellar point of interest (Fig. 3A). This alignment tool featured 20 cylindrical tracks, each a cylinder of 1/32” in diameter. All cylinders converged onto the single burr hole, from which point the trajectories diverged to arrive at the various cerebellar destinations. The 20 cylinders (guide cylinders, Fig. 3B) were configured in a 4X5 grid which corresponded to the four targeted cerebellar lobules (VIIa, VIa-c, VIe, and Vd), spanning from 0 to 2.5mm along the midline-lateral axis. The cylinders in the alignment tool provided a physical representation of the electrode trajectories above the burr hole. The tool was 3D printed using an Objet260 printer with VeroClear plastic filament.

### Alignment of the electrode

To advance the electrode, we chose a piezoelectric, high precision microdrive (0.5 micron resolution) with an integrated absolute encoder (M3-LA-3.4-15 Linear smart stage, New Scale Technologies). The microdrive held the electrode and advanced it along a single axis defined by the chosen cylinder of the electrode alignment tool. Thus, the next problem was to position the microdrive in 3D space so that its single direction of motion was precisely aligned with the desired cylinder in the electrode alignment tool.

To align the axis of the microdrive with the desired cylinder, we began by attaching the microdrive to a mechanical stereotaxic micromanipulator (SM-11-HT, Narishige, Japan). With the alignment tool installed in the base chamber, we inserted a 1/32” rod (outer diameter) inside the desired cylinder of the alignment tool (Fig. 3C), and then attached a 16G tube (0.033 inch inner diameter) to the microdrive using the same electrode holder that would also hold the electrode. The tube was fixed to the microdrive, serving as a model for the axis of travel of the electrode.

We then used the stereotaxic micromanipulator to maneuver the microdrive, thereby positioning the tube held by the electrode holder so that it precisely traveled along the axis defined by the rod. Alignment of the tube (representing the electrode) with the rod (representing the guide) was confirmed under a microscope by advancing the microdrive and demonstrating that this motion inserted the rod into the tube. Once this alignment was confirmed, the stereotactic manipulator that held the microdrive was locked into place. We then removed the tube from the microdrive and replaced it with the recording electrode (Fig. 3D). All further motion of the electrode was now under the control of the microdrive, limited to a single axis of travel that was along the desired trajectory.

### Craniotomy

Our design required that the 20 trajectories specified by the alignment tool converge on a single point on the skull and then travel within the brain to reach the cerebellum. The next step was to localize this point on the skull and drill a 1.5 mm diameter burr hole. This was achieved by using the alignment tool to mark the skull and localize the craniotomy. During the procedure the subject remained awake, following a protocol described earlier (Lu et al., 2001a).

The subject was head-fixed while the general craniotomy area, the base chamber, and the alignment tool were sterilized using alcohol, rinsed with saline, and then suctioned dry. The alignment tool was installed on the base chamber and a 22G blunt needle tip was dipped into ink and then inserted into one of the cylinders (the choice was arbitrary as all cylinders converged to the same burr hole location). The ink tipped needle was advanced until contact was made with the bone and a mark was left on the surface.

Using the procedures for aligning an electrode, we aligned a sterilized drill bit so that its trajectory of travel precisely matched the trajectory specified by a cylinder on the alignment tool. We attached a DC motor to the bit, and then attached the motor to the stereotaxic micromanipulator. At this point the alignment tool was removed, exposing the marked skull. From our co-registered pre-op and post-op CT we estimated that the total thickness of the skull and dental cement was between 900um-1100um. Under a microscope, a miniature electric drill mounted on the micromanipulator was advanced to a near touch of the dental cement. The skull and dental cement thickness estimate were used to guide the advancement of the drill bitthrough the thin layer of dental cement and about 80-90% of the thickness of the bone (before reaching the dura). Drilling concluded with observation of wet bone chips and/or liquid. The remaining bone covering the dura surface was then carefully removed using hand-held fine instruments under a microscope. Over the past two decades the Wang Laboratory (Lu et al., 2001a) has safely used this approach, and we were able to confirm that the animals never exhibited signs of discomfort or distress during this procedure.

### Sealing the burr hole with a transparent silicone gel

Our design included a single burr hole, thus requiring us to continue using a single entry for many weeks during daily electrode penetrations. To accomplish this, we experimented with a silicone gel to seal the exposed dura at the burr hole.

Silicone gels can help preserve the integrity of the intra-cranial space after a craniotomy and in prolonging the life and functionality of burr holes. We wanted to use a gel that was non-toxic, transparent, elastic, easy to apply, and had resealing capabilities, allowing multiple penetrations by electrodes over an extended period of time.

We experimented with a commercially available soft silicone gel (DOWSIL 3-4680, Dow Corning, DuraGel, Cambridge Neurotech). This gel is a polydimethylsiloxane (PDMS) based silicone and is primarily used for hydrophobic encapsulation of electronic microchips. However, due to its biocompatible and antibacterial properties, it has recently been used as a dura substitute/cover for chronic recording in mice (Jackson and Muthuswamy, 2008;Jiang et al., 2017). These previous reports had tested biocompatibility, cytotoxicity, and sealing capability of the gel in mice, suggesting that it is a safe product, effective in sealing the dura from air contact, thereby minimizing possibility of infection. The published reports suggested that the gel had a number of attractive features: it was biocompatible, reduced possibility of inflammation, reduced cerebro-spinal fluid leakage, reduced humidity loss from the craniotomy, and remained soft for a period of days to weeks, thereby allowing microelectrode penetration with minimal force.

Immediately after drilling the burr hole and cleaning the remaining bone chips we covered it with the silicone gel (DuraGel, Cambridge Neurotech). Once the gel was cured (about 30 minutes), its mechanical properties allowed the electrodes to penetrate it for weeks. Moreover, the viscosity of the gel allowed us to place it inside of the burr hole, thereby covering the dura completely, sealing it from air. The gel is transparent, which allowed us to monitor the dura beneath the gel. We replaced the gel at two week intervals.

### Data acquisition

We recorded from the cerebellum using three types of electrodes: quartz insulated 4 fiber (tetrode) or 7 fiber (heptode) metal core (platinum/tungsten 95/05) electrodes (Thomas Recording), and 64 contact high density silicon probes (Cambridge Neurotech). None of these electrodes could penetrate the marmoset dura. Therefore, we performed a micro-duratomy using a 30G needle, which was installed on the stereotaxic micromanipulator frame and advanced to the surface of dura until a puncture was made. Once the puncture was made, the various electrodes could travel through the dura.

We connected each electrode to a 32 or 64 channel headstage amplifier and digitizer (Intan Technologies, USA), and then connected the headstage to a communication system (RHD2000, Intan Technologies, USA), as shown in Fig. 4. In addition, we connected to the RHD2000 system digital outputs from our custom behavioral software and analog output from an optical sensor mounted on the TV screen.

**Fig. 4.**
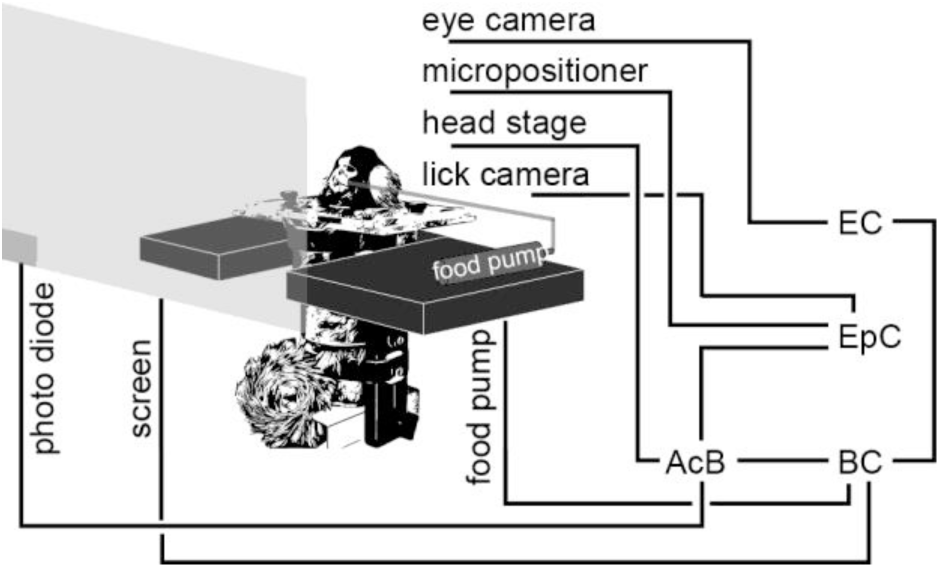
Behavioral and electrophysiological recording system. AcB: acquisition board. BC: behavioral computer. EpC: electrophysiological computer. EC: eye recording computer.

We used OpenEphys (Siegle et al., 2017), an open-source extracellular electrophysiology data acquisition software, for interfacing with the RHD2000 system and recording of signals. The signals were post processed and analyzed using MATLAB (Mathworks, USA) and Python.

### Behavioral training

In our earlier work in the macaque we had discovered that a key step in decoding activity of Purkinje cells in the oculomotor vermis was to organize them into groups in which all the cells within a group shared the same preference for visual error (Herzfeld et al., 2015;Herzfeld et al., 2018). The preference for error was signaled via the cell’s complex spike tuning, as measured when the animal made a saccade but at saccade end, the target was not on the fovea, resulting in a sensory prediction error which we represented as a vector. When the Purkinje cells were organized based on their error-dependent complex spike tuning, each population predicted in real-time the motion of the eyes during a saccade as a gain field. Thus, we trained the marmosets in a task in which we could identify the complex spike tuning of each Purkinje cell with respect to visual error.

Immediately following surgical recovery, the animal was placed on a food regulated diet. For 5 days a week, this diet consisted of 15 g of lab diet powder and 10 g of apple/mango sauce mixed in 30 g of water. This resulted in net 40 mL of deliverable food, which we administered via a syringe pump during the task. During the weekends, the food consisted of 30 g of a solid lab diet for the first day and 25 g for the second day.

Weight was monitored daily to ensure the health of the animals: weight was maintained within 85-100% of the average weight before the food regulation regime. If the weight fell below 85%, food regulation was stopped and the animal was fed in the colony until weight recovered to at least 90%. The animal was progressively acclimated to being held by a handler, entering its carrier independently, and sitting in its task chair. The experiment room was maintained at 74-84°F.

Saccade training proceeded at a frequency of 5 days/week. Visual targets were presented on a TV screen (Curved MSI 32” 144 Hz - model AG32CQ) while binocular eye movements were tracked using an EyeLink-1000 eye tracking system (SR Research, USA). We began with fixation training, followed by saccade training to a primary target. As the subject became accustomed to these paradigms, we introduced a secondary target, thus encouraging a corrective saccade (Fig. 5A).

**Fig. 5.**
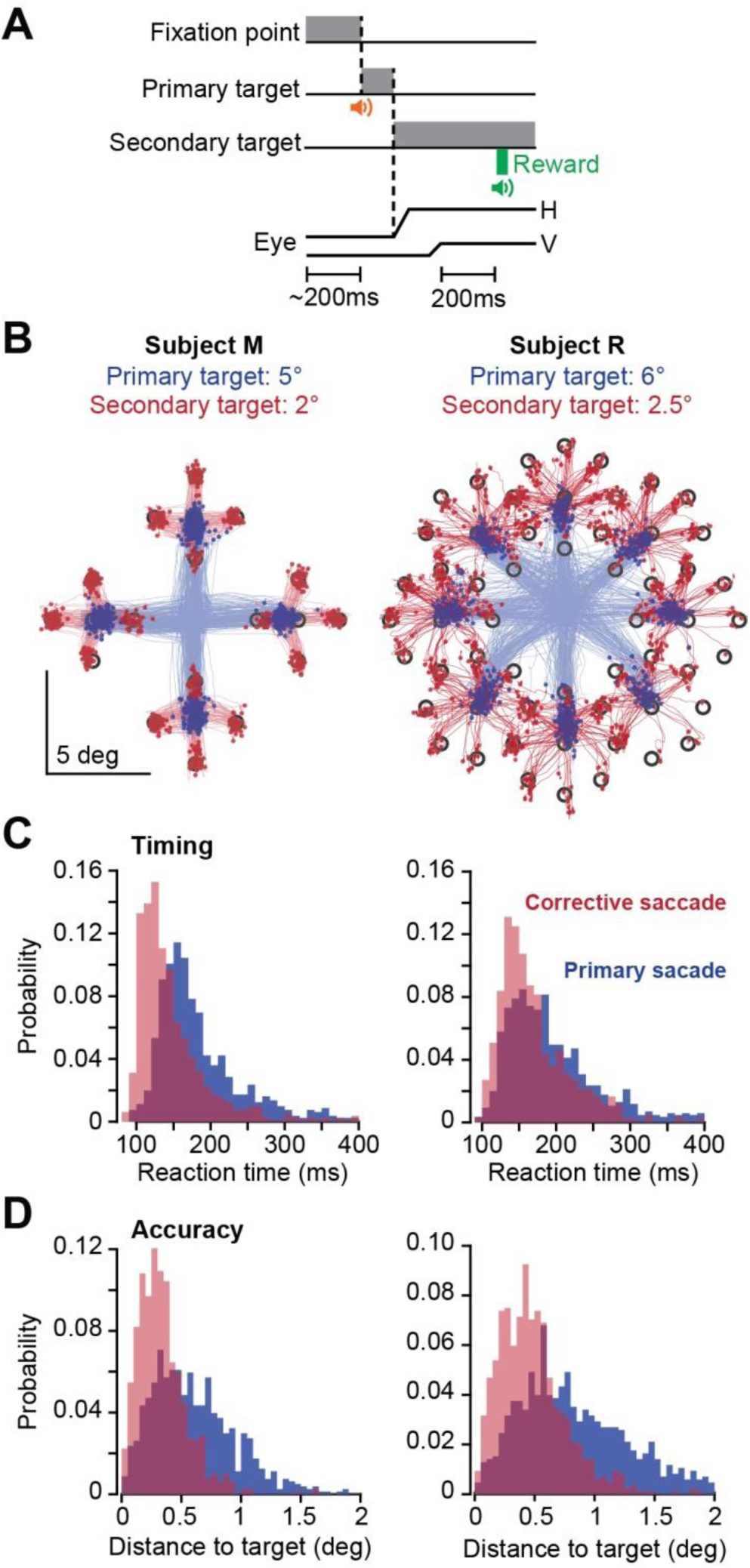
Task design and behavioral results. **A**. A trial began with 200ms of fixation, followed by presentation of a primary target at 5° or 6°. During the primary saccade the target was erased and a secondary target was presented at 2° or 2.5° displacement with respect to the primary target. Reward was presented following 200ms fixation of the secondary target. **B**. Saccade trajectories for subjects M and R during one session. Primary saccades are plotted in blue and corrective saccades are plotted in red. Targets were 0.5 ×0.5 deg square. **C**. Reaction time distribution for primary and corrective saccades. **D**. Distance to target at conclusion of primary and corrective saccades.

Following fixation training, saccade training initiated with presentation of a primary target (0.5×0.5 deg square) at a random location and distance of 3 deg. Reward was provided if the primary saccade was within 1.5° radius from the center of the primary target. In the fully trained subject the trial began with fixation of a center target for 200 ms, after which a primary target (0.5×0.5 deg square) appeared at a random location at a distance of 5-6 deg (Fig. 5A). As the animal made a saccade to this primary target, that target was erased and a secondary target was presented at a distance of 2-2.5 deg. The subject was rewarded if following the primary saccade it made a corrective saccade to the secondary target, landed within 1.5° radius of the target center, and maintained fixation for 200 ms.

Correct trials produced a distinct auditory tone, and engagement of the food pump at a rate of 0.020 mL/trial. We found that in the trained subjects, this low rate encouraged them to complete a few consecutive trials before stopping to lick the food tube, thus further increasing the number of correct trials per session.

### Data analysis

To detect simple spikes, we began with a high-pass filtered version of the signal (300 Hz), and then subtracted at each time point the mean of the signal across all contacts, a method known as common average referencing. We then used a threshold based technique to hand-sort the data.

Here, we consider only the largest spike recorded by each contact. To detect complex spikes, we relied on frequency-domain analysis that tentatively identified these spikes via their power spectrum properties.

To confirm that the complex and simple spikes originated from the same cell, we compared the conditional probability Pr(*S*(*t*)|*C*(0)) with Pr(*S*(*t*)|*S*(0)). These probabilities describe spike-triggered histograms. For example, Pr(*S*(*t*)|*C*(0)) is the probability that a simple spike occurred at time *t*, given that a complex spike was generated at time zero. Pr(*S*(*t*)|*S*(0)) is the probability that a simple spike occurred at time *t*, given that a simple spike was generated at time zero. Simple spikes that originate from a single cell produce a refractory period. Thus, Pr(*S*(*t*)|*S*(0)) should exhibit a low probability period of roughly 10 ms in duration centered at time zero. On the other hand, a complex spike suppresses production of future simple spikes, but not those that occurred before. As a result, Pr(*S*(*t*)|*C*(0)) should be asymmetric, with a long period of low simple spike probability following time point zero.

Multi-contact electrodes allow for analysis of simultaneously recorded neurons. However, spiking activity in one neuron can easily influence the data recorded by two nearby contacts, thus giving an illusion that the two contacts are picking up two distinct neurons. To guard against this, after we sorted the data in each contact, we waveform triggered the data recorded by contact A by the spikes recorded on contact B. This identified the waveform of the neuron recorded by contact B on the spike recorded on contact A. We compared this cross-contact triggered waveform with the within contact triggered waveform generated by the spikes recorded by contact A. The cross-contact triggered waveform should produce a different cluster of spikes in A than the main neuron isolated by A. If there were spikes that occurred within 1 ms of each other on contacts A and B, we used these coincident-spike events to trigger the waveform in A. The spikes in A that were identified to be coincident with B should look approximately the same as the non-coincident spikes in A.

To quantify coordination between activities of two Purkinje cells, we computed conditional probabilities. For contacts 1 and 2, we computed Pr(*S*1(*t*)|*C*2(0)), testing whether complex spikes on contact 2 produced any changes in the simple spikes on contact 1. If the data were sorted properly, this conditional probability should be essentially flat. To visualize coordination between the neurons, we next plotted the spike-triggered waveform of voltages recorded by contact 1, triggered by the simple spikes on contact 2. Finally, we computed Pr(*S*1(*t*)|*S*2(0)). This measure quantified whether the occurrence of a simple spike on contact 2 altered the probability of simple spikes on contact 1.

## Results

The methods that we described above allowed us to train subjects to produce around a thousand trials per session during cerebellar recordings. However, we arrived at these methods following a learning process that involved a number of unsuccessful attempts. Here, we describe our experience in terms of both types of results.

### Behavioral results

For subject B, we experimented with saccade training without the head-post using a loose fitting head-restraining system (days -416 to 0, training without head-post, Fig. 6A). Following a number of attempts we abandoned this approach because we were unable to produce robust calibration.

**Fig. 6.**
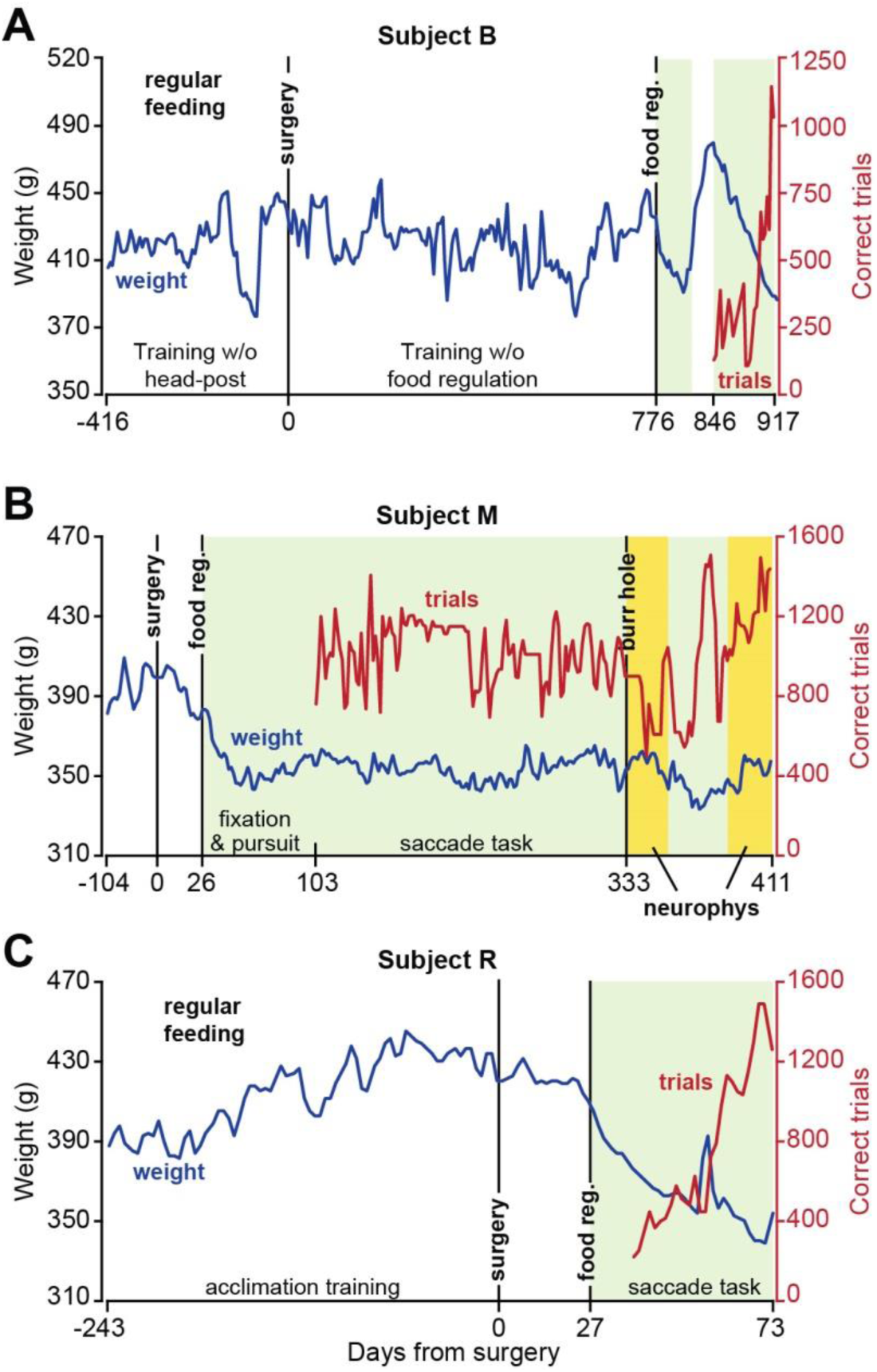
Weight patterns and number of correct trials following food regulation. **A**. Record of weight and correct trials in subject B. Training began without a head-post, but was unsuccessful because of an inability to produce reliable calibration of eye measurements. Following surgery, a long period of training without food regulation ensued, but was ultimately abandoned because of the unwillingness of the animal to work for more than a few hundred trials. Following food regulation, performance in the task dramatically improved. **B**. Weight and correct trial data in subject M. **C**. Weight and correct trial data in subject R. Correct trials data represent running average of bin size 2. Green color indicates periods of food regulation. Yellow color indicates periods of food regulation and neurophysiological recording.

We began behavioral training of subject B after head-post surgery using the traditional method (Mitchell et al., 2014;Johnston et al., 2018) of feeding the animals in the colony and supplementing that food with reward during task performance in the head-fixed condition (training without food regulation, Fig. 6A). We found that this was insufficient to motivate this subject.

We next eliminated home-cage feeding on experiment days, and instead provided task-based food rewards during the head-fixation period (food regulation, Fig. 6). We ensured that regardless of performance, the subjects always received their required allotment of food (at least 25 ml of food per day) and weight was maintained at around 90% of the pre-food regulation regime.

However, food was delivered only while the animal was head-fixed. This produced improved performance in subject B (days 846-917, Fig. 6A). We then fully tested this approach in subjects M and R (Fig. 6B and Fig. 6C).

Following recovery from surgery, subject M was food regulated and fed only in the head-fixed condition. After 30 sessions of fixation training, subject M was trained for 20 sessions on trials with primary saccades only (gradually increasing from 3° to 5°). Following this, subject M began training on the main task (Fig. 5A), which included both primary and corrective saccades. The correct number of trials shown in Fig. 6B begins with the first day of training on the primary saccade, and continues as the subject transitioned to the main task. We found that within three months after start of fixation training subject M was able to perform around 1000 correct trials in the main task, and then maintained this performance for over a year. During neurophysiological experiments, subject M produced 600-1500 correct trials per session (neurophysiology trials, running average data presented in Fig. 6B).

We experimented with a different approach to saccade training in subject R. Instead of starting with fixation training, we began with pursuit training for 3 days. This subject’s good performance allowed us to subsequently train on a saccade task for 15 sessions during which only the primary target was presented and amplitude gradually increased from 3° to 5°. The subject was then trained on the main task (Fig. 5A), during which the primary target was placed at 6° and the secondary target was at 2.5°. The correct number of trials in Fig. 6C begins with the first day of training on the primary saccade, and continues as the subject graduated to the main task. This subject was able to produce 1000-1500 correct trials in the main task within two months after food regulation (Fig. 6C).

Notably, during a year of food-regulated training in subject M, and two months of food-regulated training in subject R, both remained healthy, as suggested by their stable weight and lack of complications.

Our aim was to train the subjects in a task in which we could measure contributions of Purkinje cells to control of saccades. Each trial began with 200 ms fixation of a central location, and was followed by a primary saccade to a peripheral target (0.5×0.5 deg square) at 5° or 6° displacement, corrective saccade to a secondary target at 2° or 2.5° displacement, and 200 ms of fixation at the secondary target (Fig. 5A). Saccades for a representative session are plotted in Fig. 5B, and the distribution of reaction times are plotted in Fig. 5C. Primary saccade reaction times for subjects M and R were 188±65 ms (mean±SD) and 197±69 ms. Corrective saccade reaction times for subjects M and R were 146±50 ms and 169±50 ms. To quantify saccade accuracy, we measured the distance between saccade endpoint and target center and found that for the primary saccade, this distance was 0.59±0.35 deg (mean±SD) for subject M, and 0.85±0.50 deg for subject R (Fig. 5D). For the corrective saccade, distance to target was 0.34±0.22 deg for subject M and 0.49±0.32 deg for subject R. The reward region was 1.5° radius around the target. As a result, across the two subjects 81% (1011 out of 1251) and 88% (1464 out of 1672) of the trials that initiated with a primary saccade to the target concluded correctly and were rewarded.

An important component of our training was that the subjects received a small amount of food per correct trial: 0.02 mL per trial. At completion of a successful trial, the computer generated a beep, and food delivery was signaled by the sound of the pump. However, the low reward rate encouraged the subjects to withhold licking until the food accumulated in the transparent tube that was placed in front of them. The effect was to teach the subjects to perform the task in blocks of 3-4 uninterrupted consecutive trials, thus increasing the total number of successful trials per session.

### Simultaneous recordings from pairs of Purkinje cells

We explored lobules V, VI and VII of the cerebellum using tetrodes, heptodes, and high density silicon arrays. We thought that the cranial ground screw would be required to provide a reference for our electrophysiological recordings. However, with experience with subject M we learned that the ground screw was unnecessary: the head-post by itself acted as a reliable reference for the electrical signals during neurophysiological recordings. This is likely because the titanium screws established electrical continuity between the head-post and the bone. Thus, in subject R we eliminated the ground screw from the surgical procedures.

As the electrode approached the tentorium, there was a noticeable increase in background activity, indicative of proximity to Purkinje cells and their high baseline discharge rate. When the electrode passed through the tentorium, there was a distinct “pop”, which was then followed by a gradual increase in spiking activity.

Identification of a Purkinje cell required presence of both simple and complex spikes. Examples of putative complex and simple spikes, recorded by a silicon array, a tetrode, and a heptode are presented in Fig. 7A. Complex spikes have a stereotypical positive wave that has slow dynamics, resulting in greater power in the lower frequencies (<300 Hz range) than the waveform of simple spikes (Warnaar et al., 2015). Furthermore, complex spikes are relatively rare events, presenting an inter-spike interval that is roughly two orders of magnitude greater than simple spikes.

**Fig. 7.**
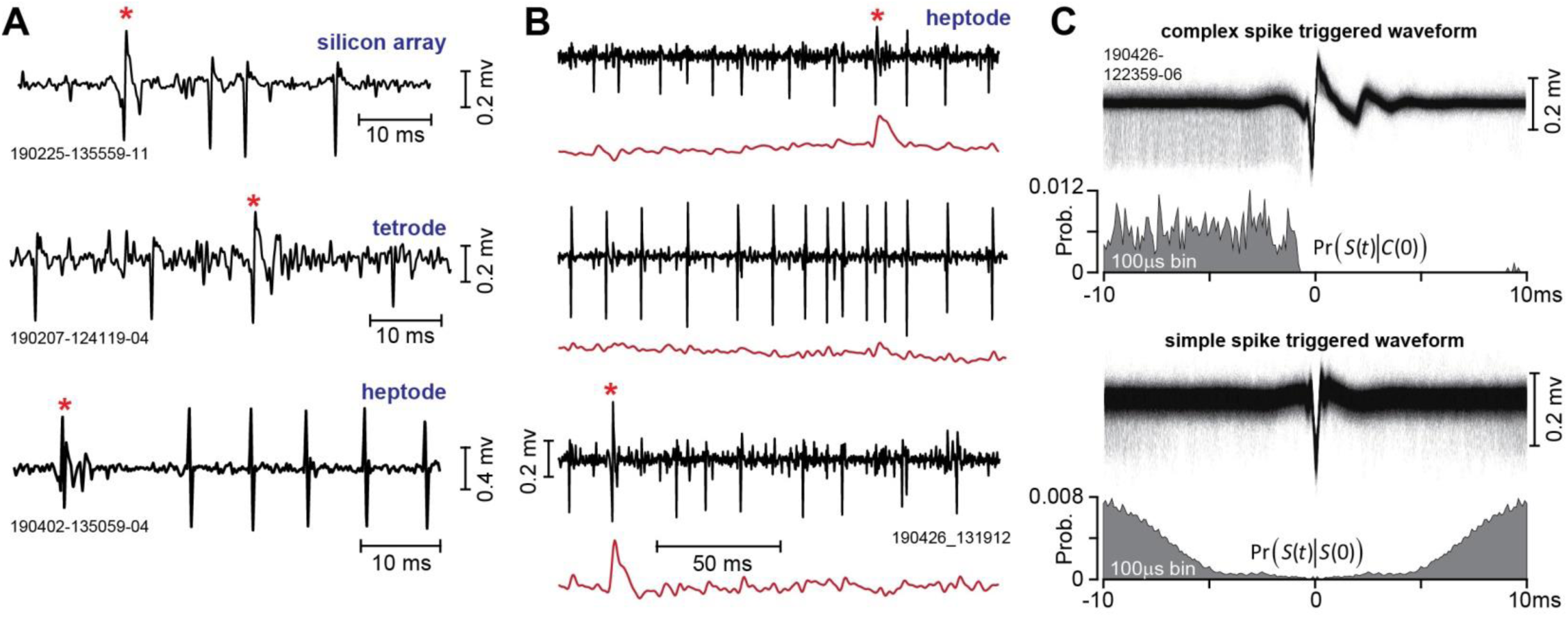
Recordings from the marmoset cerebellum. **A**. Complex and simple spikes recorded by a silicon array, a tetrode, and a heptode. The complex spike is noted by an asterisk. **B**. Example of multiple neurons simultaneously recorded by a heptode. The black traces are the high-pass filtered voltages and the red trace is the low-pass filtered version of the same signal (300Hz cutoff). The top and bottom traces are putative Purkinje cells while the middle trace is an unidentified third neuron. **C**. Spike-triggered waveforms and histograms. In the upper trace, the voltages recorded by a single contact on a heptode were triggered by the complex spikes on the same contact. The resulting complex spike-triggered waveform exhibits a uniform pattern of simple spikes before the complex spike, and then simple spike suppression. This pattern is quantified in Pr(*S*(*t*)|*C*(0)), which is the probability of a simple spike at time *t*, given that a complex spike occurred at time 0. In the lower trace, the voltages were triggered by the simple spikes. The resulting simple spike-triggered waveform exhibits a symmetric period of spike suppression, reflecting the Purkinje cell’s refractory period. This pattern is quantified in Pr(*S*(*t*)|*S*(0)).

A useful real-time tool for identification of complex spikes is a low-pass filter (300 Hz), which under good recording conditions can provide a tentative label for the complex spikes. Fig. 7B shows an example of recording data from a heptode that simultaneously isolated three neurons: two putative Purkinje cells, and a third unidentified neuron. In this figure, the voltages for each of the three simultaneously recorded neurons are represented as black traces, and the low-pass filtered version of the same signal is shown in red. The complex spikes are tentatively identified by the sharp increase in the low-frequency power of the signal.

An important feature of a Purkinje cell is that production of a complex spike suppresses the generation of simple spikes for a period of 10ms or longer. Thus, the shared cellular origin of complex and simple spikes can be verified by using the complex spikes to trigger the voltage waveform. The upper trace of Fig. 7C provides an example of this, demonstrating that the simple spikes occur with roughly equal probability before the complex spike, but then cease entirely after the complex spike.

We quantified this pattern via the condition probability Pr(*S*(*t*)|*C*(0)), which measured the probability of a simple spike at time *t*, given that a complex spike occurred at time 0. This measure illustrated that the simple spikes occurred with equal frequency before the complex spike, but were then completely suppressed for around 10ms following the complex spike.

A second feature of a Purkinje cell is that the simple spikes have a short refractory period. This feature can be verified by using the simple spikes to trigger the voltage waveform. The lower trace of Fig. 7C provides an example of this. We quantified this pattern via the conditional probability Pr(*S*(*t*)|*S*(0)), which measured the probability of a simple spike at time *t*, given that a simple spike occurred at time 0. This measure illustrated that the simple spikes exhibited a 5ms refractory period.

Two examples of simultaneously recorded neurons from lobule VII during the saccade task are shown in Fig. 8. In Fig. 8A, the heptode isolated three neurons, two of which were Purkinje cells (high-pass filtered signals are shown in black, the low-pass filtered signal is shown in red). In this trial, the animal was presented with a target at 5° on the horizontal axis and made a saccade at 220ms latency (bottom row, Fig. 8A). At saccade onset the target was erased and redrawn at a displacement of 2° along the vertical axis. Thus, at the conclusion of the vertical saccade the target was not on the fovea, generating a visual error, and encouraging the animal to make a corrective vertical saccade. This is an example of cross-axis adaptation (Deubel, 1987;Xu-Wilson et al., 2009). The saccade landed about 0.5° short of the target, and was followed by a corrective saccade. The Purkinje cell on contact 6 (2nd row, Fig. 8A) produced a complex spike before the primary saccade. A second Purkinje cell, recorded on contact 5 (top row, Fig. 8A), did not produce a complex spike before the onset of the primary saccade, but rather produced a complex spike following experience of a visual error (before onset of the corrective saccade). A third neuron was isolated by contact 7, but this neuron was not a Purkinje cell as it lacked complex spikes.

**Fig. 8.**
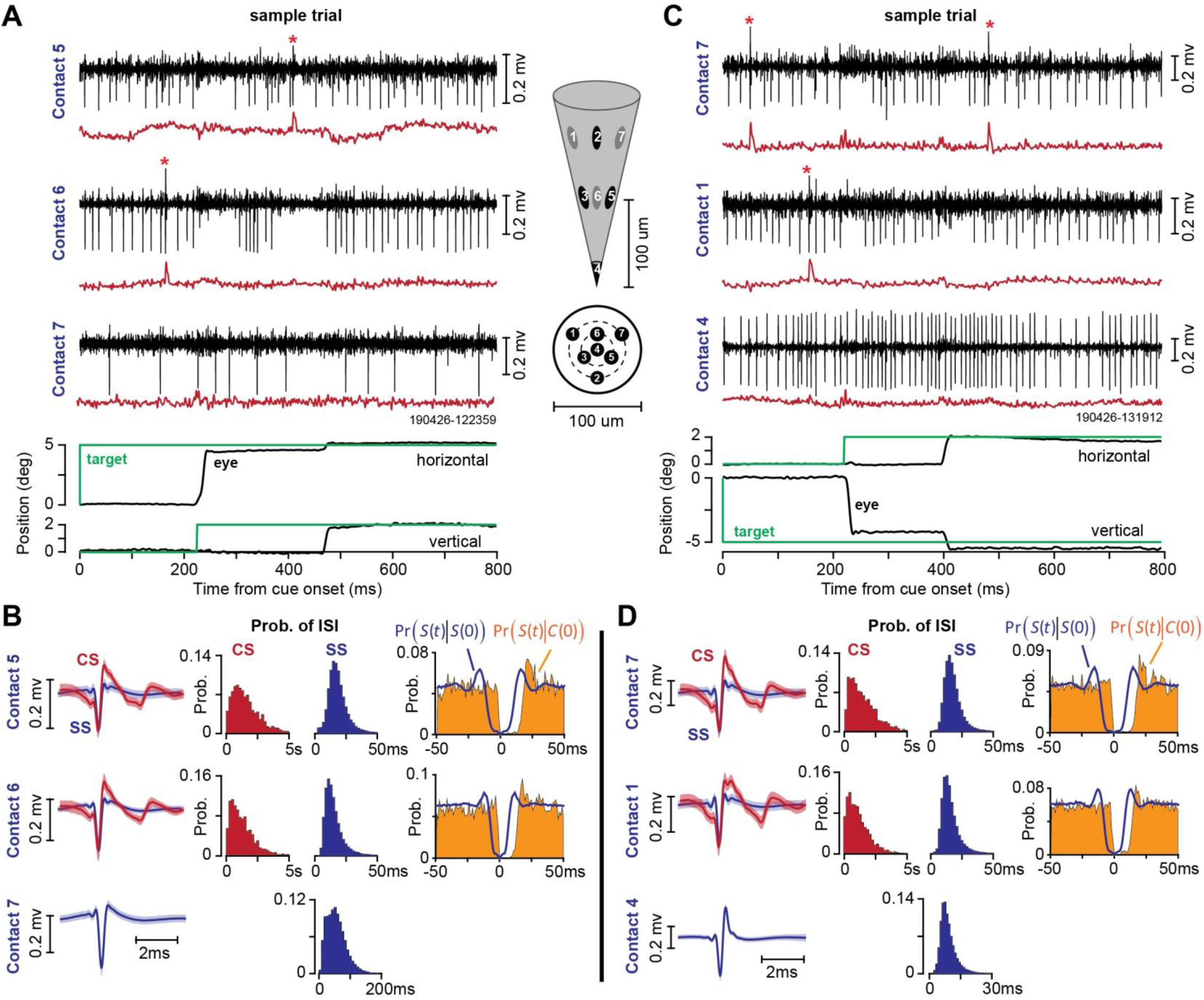
Example of heptode recordings during two sessions in the saccade task. **A**. Sample trial, illustrating activity of a Purkinje cell on contact 5, another Purkinje cell on contact 6, and an unidentified neuron on contact 7. The black traces are high-pass filtered voltages and the red traces are low-passed filtered signal (300 Hz cutoff). The middle figure shows the geometry of the heptode and the location of each contact. **B**. Spike waveforms and patterns of spike timing recorded from the session in part A. CS is complex spike and SS is simple spike. ISI is inter-spike interval. Pr(*S*(*t*)|*S*(0)) is the probability of a simple spike at time *t*, given that a simple spike occurred at time zero. Pr(*S*(*t*)|*C*(0)) is the probability of a simple spike at time *t*, given that a complex spike occurred at time zero. **C**. Sample trial recorded in a different session, illustrating activity of a Purkinje cell on contact 7, another Purkinje cell on contact 1, and an unidentified neuron on contact 4. **D**. Spike waveforms and patterns of spike timing recorded from the session in part B. Bin size is 1 ms for conditional probabilities, 200 ms for probability of complex spike ISI, and 2 ms for probability of simple spike ISI. Error bars on the spike waveforms are standard deviation.

The waveforms for the complex and simple spikes in this recording session for each contact are shown in the left column of Fig. 8B. The waveforms illustrate the slower dynamics of the complex spikes, thus providing the basis for why a low pass filter is generally useful in their identification.

The middle column of Fig. 8B displays the probability densities of the inter-spike intervals for each type of spike (note that the x-axis for the complex spikes is 5 seconds, whereas for the simple spikes is 50 msec). The number of simple spikes in a given period of time was roughly two orders of magnitude larger than the number of complex spikes.

The right column of Fig. 8B illustrates the within-cell timing properties of the spikes via conditional probabilities. Pr(*S*(*t*)|*C*(0)) demonstrates that following a complex spike at time 0, there was a silent period of ∼10 ms during which simple spikes were absent. Thus, production of a complex spike briefly suppressed production of simple spikes, demonstrating that the complex and simple spikes on this contact originated from the same Purkinje cell. Pr(*S*(*t*)|*S*(0)) demonstrates a 5 ms refractory period, which is critical for demonstrating that the simple spikes were correctly attributed to a single Purkinje cell, and not misattributed because of cross-contaminated spikes from a neighboring contact. Finally, a comparison of Pr(*S*(*t*)|*C*(0)) with Pr(*S*(*t*)|*S*(0)) demonstrates that the complex-spike induced suppression of simple spikes was longer than the typical simple spike refractory period, another indication that the source of complex and simple spikes was a single Purkinje cells.

Another example of a recording session is shown in Fig. 8C. In this trial, the target was presented at 5° along the vertical axis, and the animal made a saccade at a latency of 220 ms. At saccade onset, the primary target was erased and a new target at 2° to the right of the primary target was displayed. A Purkinje cell was isolated on contact 1, while a second Purkinje cell was isolated on contact 7. Contact 4 isolated a third neuron, but this was not a Purkinje cell.

Fig. 8D illustrates the within contact timing properties of the complex and simple spikes. Like the data in Fig. 8B, the conditional probabilities demonstrate that simple spikes exhibited a 5 ms refractory period, and following a complex spike, there was suppression of simple spikes for a period of 10 ms.

In our experience, the head-post and the electrode holder provided good recording stability. Over the course of 22 sessions in which single or pairs of Purkinje cells were isolated, we were able to maintain single unit isolation during an average of 39.1±3.8 minutes (mean±SEM), producing 516±62 correct trials.

Activity of a Purkinje cell over the course of a representative session (700 trials, 40 minutes) is illustrated in Fig. 9. In this session, subject M performed the task shown in Fig. 5B. The trial began with a primary target at 5° displacement. Upon saccade initiation, the primary target was erased and replaced with a secondary target at 2° displacement (Fig. 9A). A corrective saccade followed termination of the primary saccade at a latency of 120-150 ms. The rasters in Fig. 9A present the timing of simple and complex spikes during each trial via thin blue and thick red lines, respectively. This cell produced a burst of spikes around primary saccade onset, exhibiting elevated activity that continued long after the saccade had ended (Fig. 9B, left panel). There was a second burst of simple spikes around the time of the corrective saccade (Fig. 9B, right panel), which also exhibited a duration that was much longer than the saccade. This disparity between burst duration of simple spikes and duration of saccade is a common feature of saccade related Purkinje cells (Thier et al., 2000;Herzfeld et al., 2015).

**Fig. 9.**
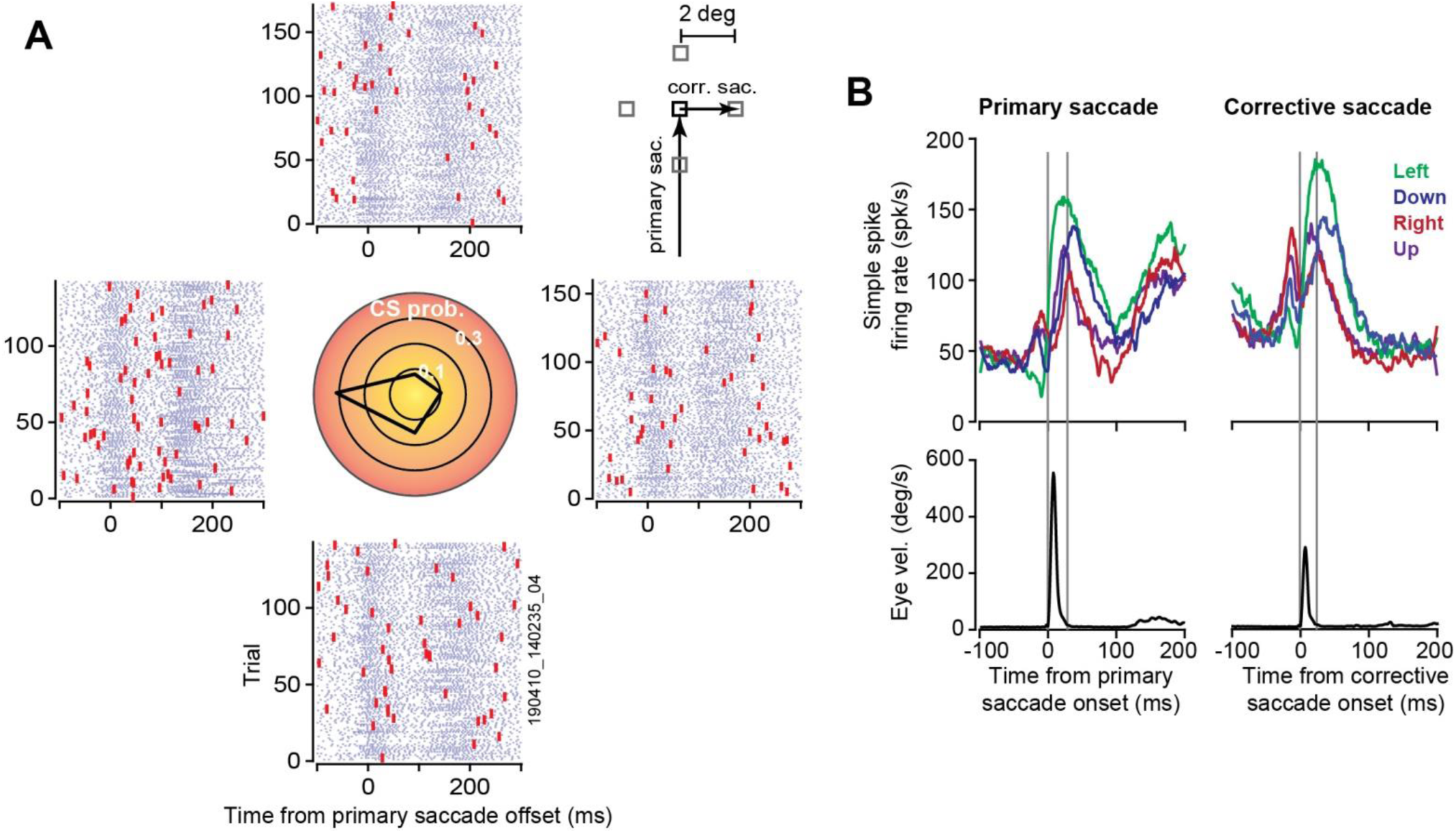
Example of data recorded from a Purkinje cell (subject M) during the task shown in Fig. 5B. **A**. The primary target (black square) was presented at 5° displacement. Upon initiation of saccade, the target was replaced with a secondary target (gray square) at 2° displacement in one of four directions, inducing a visual error and encouraging a corrective saccade. Rasters are aligned to primary saccade offset, and grouped based on the direction of the visual error vector. For example, the rasters on the right refer to trials in which the corrective saccade was to the right. The central plot shows probability of complex spikes during the 200ms period following offset of the primary saccade. This cell had a preference for visual errors toward -180°. **B**. Simple spike activity and eye velocity aligned to onset of primary saccade and corrective saccade. In the left subplot the simple spikes are grouped based on direction of the primary saccade. In the right subplot the spikes are organized based on direction of the corrective saccade. The gray vertical lines denote onset and offset of the saccade. The cell exhibited a burst of activity around saccade onset, but burst duration was much longer than saccade duration.

Upon termination of the primary saccade the target was not on the fovea, resulting in a sensory prediction error. This visual event likely engaged neurons in the superior colliculus (Kojima and Soetedjo, 2017) and the inferior olive, leading to occasional complex spikes. We measured probability of complex spikes during the 200 ms period following completion of the primary saccade (central panel of Fig. 9A). This Purkinje cell was tuned to visual errors along -180°, suggesting that it received visual error information from the right superior colliculus via the left inferior olive.

Our earlier work (Herzfeld et al., 2015;Herzfeld et al., 2018) found that when Purkinje cells are organized into populations that share a common preference for error (i.e., complex spike tuning is similar), their combined simple spike activity produces a pattern of spikes that no longer has a duration disparity with respect to the saccade. That is, while the simple spike activity of individual Purkinje cells does not have an obvious relationship to eye motion during a saccade, as a population the Purkinje cells can precisely predict motion of the eye in real-time.

### Spike timing properties of simultaneously recorded Purkinje cells

The ability to record simultaneously from multiple Purkinje cells allows one to ask whether nearby cells coordinate their activity, a function that may be fundamental in driving neurons in the cerebellar nucleus (Person and Raman, 2012). However, with multiple contacts there is a danger that some of the spikes that are recorded on contact A and attributed to Purkinje cell 1 are in fact generated by nearby Purkinje cell 2, which in turn is also being recorded by contact B. This cross-contact influence (cell 2 affecting recordings in both contacts A and B) would result in spurious coordination. One way to guard against this misattribution is to compare the spike-triggered waveforms.

Fig. 10A (left column) illustrates the waveform of simple spikes recorded by contact 6, during the same session as that shown in Fig. 8A. The middle column illustrates the waveform recorded by contact 6 but triggered by the simple spikes on contact 5. The data suggest that on average, the voltages recorded by contact 6 reflect in only a minor way the spikes being recorded by contact 5. Occasionally, the two P-cells produced a simple spike within 1ms of each other. The waveform on contact 6 as triggered by only these synchronous events (right column of Fig. 10A) looks very similar to the waveform of the usual spikes (left column of Fig. 10A). Thus, these features suggest that any coordinated activity present in the data of these two contacts would be because two independent cells fired simple spikes at approximately the same time.

**Fig. 10.**
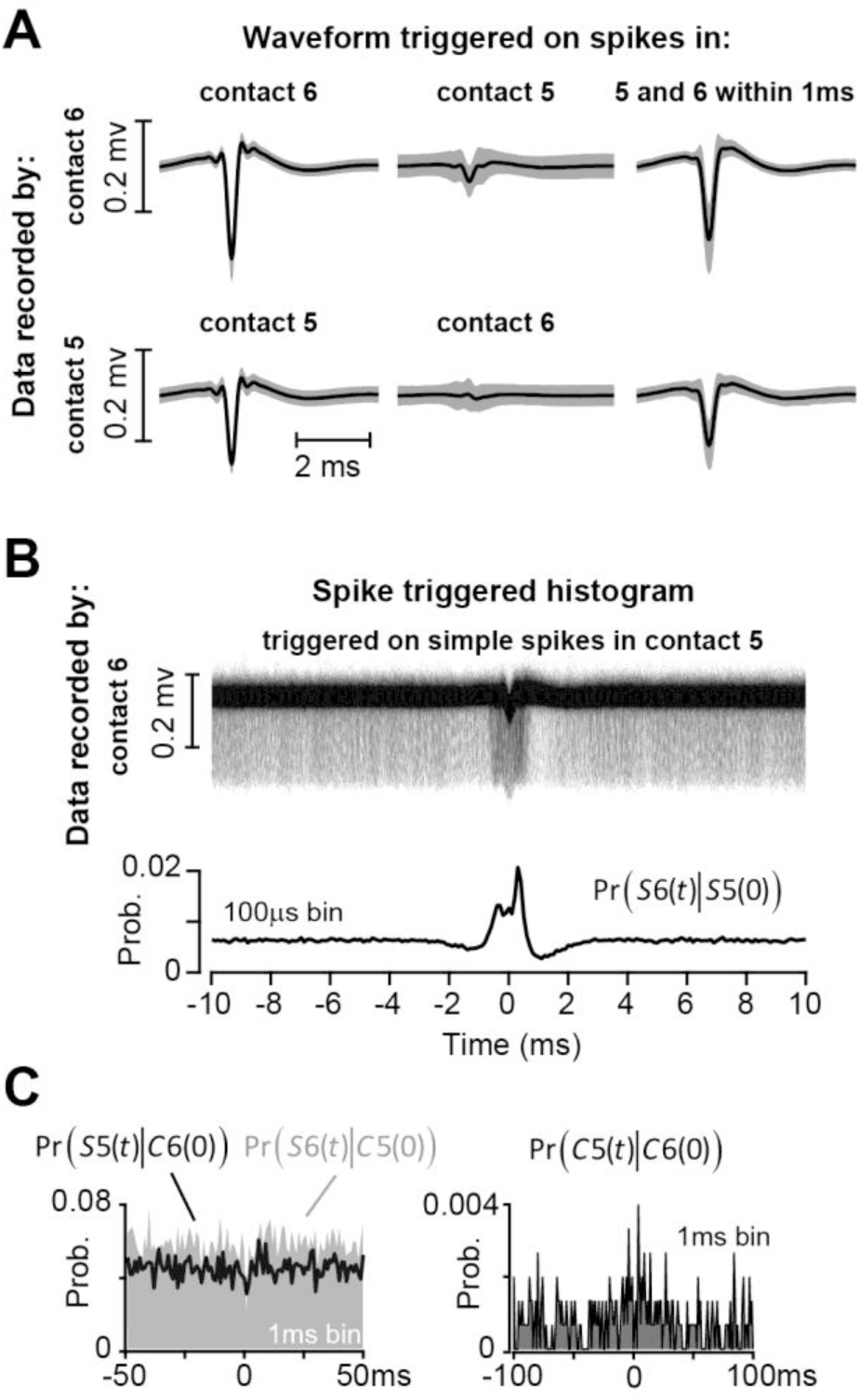
Spike timing property of a pair of simultaneously recorded Purkinje cells during the saccade task. **A**. A Purkinje cell was isolated by contact 5, and another one by contact 6. Upper row: data recorded by contact 6 were triggered by simple spike events on contact 6 (left column), and simple spike events on contact 5 (middle column). The right column is data recorded by contact 6 and triggered if there was a simple spike event in both contacts 6 and 5 within 1ms of each other. Lower row shows the waveform for contact 5. On average, spikes in one contact did not produce a significant voltage change in another contact, and the shapes of synchronous (within 1ms) and non-synchronous spikes were nearly identical. **B**. To measure coordination among the Purkinje cells, we triggered the waveform on contact 6 by the spikes on contact 5. The pattern of coordination is quantified via the conditional probability Pr(*S*6(*t*)|*S*5(0)). **C**. Left plot shows the probability of simple spikes at time *t*, given that a complex spike occurred at time zero in another channel. There is no suppression of simple spikes following a complex spike. The plot on the right shows the probability of co-occurrence of the complex spikes. The two cells did not produce complex spikes that were strongly coordinated.

We visualized the coordination between two Purkinje cells by triggering the voltages recorded by one contact via the simple spikes generated by a different Purkinje cell on a separate contact (Han et al., 2018). The resulting spike-triggered histogram of waveforms recorded by contact 6, triggered by spikes on contact 5, is shown in Fig. 10B. The term Pr(*S*6(*t*)|*S*5(0)) measures the probability of simple spikes on contact 6 at time *t*, given that a simple spike occurred on contact 5 at time 0. The conditional probability exhibits a peak at around 0ms, suggesting a measure of coordination. This coordination could have arisen because both Purkinje cells were saccade related, or driven by presentation of a visual stimulus. It is also possible that these two cells received similar inputs (Heck et al., 2007), or that activity in one affected activity in the other (Han et al., 2018).

The left plot of Fig. 10C illustrates the spike timing properties of these pairs of Purkinje cells. The conditional probability Pr(*S*5(*t*)|*C*6(0)) quantifies the effect of complex spikes on contact 6 at time 0 on the simple spikes on contact 5 at time *t*. It demonstrates that the complex spike on contact 6 did not suppress the simple spikes on contact 5. Similarly, the conditional probability Pr(*S*6(*t*)|*C*5(0)) demonstrates that the complex spike on contact 5 did not suppress the simple spikes on contact 6. This provides further evidence that the two nearby contacts picked up two distinct Purkinje cells.

The conditional probability Pr(*C*5(*t*)|*C*6(0)), displayed in the right plot of Fig. 10C, quantifies the relationship between complex spikes on contacts 5 and 6. This plot illustrates that the complex spikes in these two Purkinje cells were not well coordinated (little or no co-occurrence at 0ms latency).

In summary, these data from a pair of simultaneously recorded Purkinje cells, cells that were likely less than 50 μm apart, suggested that the cells were not served by the same inferior olive neuron. However, the simple spikes exhibited a measure of coordination, suggesting that they may share common inputs, or that the electrical activity in one cell influenced the activity in the other (Han et al., 2018).

### Silicone gel coating of the burr hole

We sealed the burr hole with a silicone based polymer (DuraGel) that we hoped would provide long-term protection to the dura, while allowing for daily penetrations. Our experience with this product has been positive. Fig. 11 provides images of the burr hole after drilling and following application of the gel. The gel remained in place for two weeks as we performed daily recordings. At two week intervals we removed the gel, examined the dura under a microscope, and took samples and performed tissue culture. In every case the results suggested a healthy dura without any evidence of infection. Thus, DuraGel appears compatible with the marmoset dura, providing long-term protection while allowing for daily electrode penetration.

**Fig. 11.**
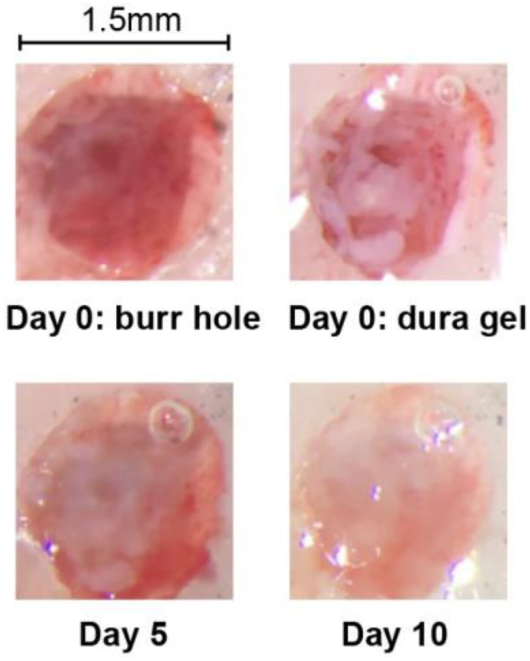
Images from the burr hole before (day 0) and after application of the silicone gel.

## Discussion

While the marmoset presents a number of attractive features as a primate model for the study of neural control of social behaviors, verbal communication, and saccadic eye movements, there is concern regarding whether the animals can be motivated to produce a sufficient number of trials. Previous reports on head-restrained marmosets have demonstrated operant conditioned behaviors in auditory tasks (Osmanski and Wang, 2011;Osmanski et al., 2013;Remington et al., 2012) as well as eye movement tasks (Mitchell et al., 2014;Johnston et al., 2018). However, performance has been a concern: around 100 trials per session in the auditory task (Osmanski and Wang, 2011;Osmanski et al., 2013;Remington et al., 2012), 80-100 trials per session in the saccade task (Johnston et al., 2018), and 300-800 trials per session in the eye movement task (Mitchell et al., 2014). Here, we aimed to improve on these results, and found that through a behavioral training protocol, it was possible to motivate the animals to consistently perform around 1000 trials per session.

In addition, in order to investigate the neural basis of motor learning, we aimed to build tools that allowed for neurophysiological investigation of the cerebellum. We began with a CT+MRI based geometric model of the subject’s skull, and used that data to compute the curvature of the base of a 3D printed titanium head-post and chamber. Using the imaging data, we designed an alignment tool that had cylinders that guided an electrode through the burr hole via an absolute encoder microdrive. This resulted in stable recordings from multiple Purkinje cells during the saccade task.

### Behavioral training

In previous reports, marmosets typically underwent a moderate food regulation regime and then were presented with the opportunity to acquire a form of liquid reward such as diluted apple sauce. In our approach, the subjects were gradually trained to receive their main source of nutrition as a consequence of rewarded behavior. Although our experience is limited to only three subjects, the results are encouraging: we were able to maintain the subjects in excellent health, while motivating them to perform the task.

In previous reports, the amount of reward delivered per trial was relatively large: 0.1-0.2 mL per trial (Osmanski and Wang, 2011;Osmanski et al., 2013;Remington et al., 2012), 0.07 mL per trial (Johnston et al., 2018), or 0.05-0.06 mL per trial (Mitchell et al., 2014). Here, we trained the subjects to expect a lower reward rate, 0.02 mL per trial. The sound of the successful trial and the engagement of the food pump indicated food delivery, but the low reward rate encouraged the subjects to wait three or more correct trials before harvesting the accumulated food. This low reward rate may have been an important factor in motivating the subjects to consistently produce a relatively large number of trials per session.

We were able to substantially reduce the training period in subject R by eliminating fixation training and instead starting with pursuit, finding that after only a few days it was possible to begin training on the saccade task, and arrive at around a thousand correct trials within 1.5 months after start of food regulation.

### Head-post design

Our head-post design departed significantly from epoxy-based helmet design that was employed in the past two decades in the study of the marmoset auditory cortex (Lu et al., 2001b;Wang et al., 2005;Eliades and Wang, 2008;Roy and Wang, 2012). Our design was made possible because of recent advances in 3D printing of titanium, allowing the use of imaging to build a subject-specific head-post.

The X-shaped design of our head-post was in contrast to the halo design recently demonstrated by the Everling group (Johnston et al., 2018) in the marmoset. Each design has its own advantages. The halo design maximizes the open surface of the skull, thus allowing for access to many regions of the brain, which is ideal for simultaneous recording from multiple cortical areas. In contrast, our X-design limits the access to only the posterior regions, but has the advantage that the animal’s skin can cover the entire head-post base, leaving only the chamber and the head-post pole exposed. Thus, our design allowed the skull to remain largely covered by skin.

### Simultaneous recording from multiple Purkinje cells in the marmoset cerebellum

Although the marmoset is roughly the same weight as a rat, its cerebellum is about twice the size (Fujita et al., 2010). Similar to the rat, stripes of different aldolase C expression intensities separate the marmoset cerebellum into longitudinal compartments. However, olivo-cortical and cortico-nuclear projections indicate that the marmoset cerebellum has several compartments that are not present in rodents (compartments in the flocculus, nodulus, and the most lateral hemispheres) (Fujita et al., 2010). In comparison to the macaque, marmoset cerebellum has fewer folia, but a relatively larger vermis. Because the vermis is likely the region critical for saccadic eye movements, as well as control of the tongue during vocalization, the marmoset vermis may be particularly well suited for study of internal models in motor control.

We found that with modern electrodes it was possible to isolate multiple Purkinje cells from the vermis during goal-directed saccades, something that to our knowledge has only once before been reported in the primate (Medina and Lisberger, 2007). We have experimented with tetrodes, heptodes, and high density silicon arrays, finding that each has its own advantages. Tetrodes and heptodes have a 3D geometry that give them the potential to record from multiple Purkinje cells within the same folium, whereas silicon arrays provide the possibility to record from Purkinje cells across folia. In our hands, the 3D geometry of the heptode has provided a robust ability to routinely isolate and stably maintain recording from pairs of Purkinje cells. These cells often share important functional properties such as complex spike tuning and simple spike coordination (Fig. 10).

We also found that the computational tools used for identifying Purkinje cells in other animals worked well in the marmoset. Like other animals, the complex spikes in the marmoset Purkinje cells had a waveform that exhibited slow dynamics, making it possible to use frequency based filtering techniques to tentatively distinguish complex spikes from simple spikes. Furthermore, following a complex spike, there was a 10-15ms period of simple spike suppression, helping to confirm that a single Purkinje cell produced both spikes.

Finally, we were able to simultaneously record from pairs of Purkinje cells and observe sub-millisecond coordination in their timing. Populations of Purkinje cells can coordinate their spiking, and this coordination may play a critical role in regulating activity of neurons in the deep cerebellar nucleus (Person and Raman, 2012). Perhaps the earliest demonstration of simple spike coordination was by Bell and Grimm (1969), who made microelectrode recordings in anesthetized cats, finding that Purkinje cells that were located <70um apart often fired nearly simultaneously. Later work found further evidence in support of this idea (Ebner and Bloedel, 1981;De Zeeuw et al., 1997;Shin and De, 2006;Heck et al., 2007;Wise et al., 2010). For example, Heck et al. (2007) demonstrated that in rats, P-cells within a few hundred microns fired synchronously during the period of reach to grasp of a food pellet, but not during the period after grasp completion.

Here we observed some evidence for sub-millisecond coordination in simple spikes of nearby Purkinje cells during performance of a saccade task. A recent study in mice (Han et al., 2018) also found that firing of many neighboring Purkinje cells exhibited sub-millisecond coordination. That study noted that when chemical synapses were blocked in brain slices, some synchrony persisted. They suggested that sub-millisecond coordination of neighboring Purkinje cells was not via shared synaptic inputs, but via electrical activity that in one Purkinje cell resulted in opening of sodium channels in the neighboring Purkinje cell. The question of whether Purkinje cells in the primate produce coordinated activity during goal-directed movements remains to be explored.

### The need to record from Purkinje cells during sensorimotor learning

In order to move accurately, the brain relies on the cerebellum to build internal models and predict sensory consequences of motor commands (Maschke et al., 2004;Smith and Shadmehr, 2005;Izawa et al., 2012;Golla et al., 2008;Rabe et al., 2009;Donchin et al., 2012;Roth et al., 2013). However, it has been difficult to decipher how the cerebellum learns internal models because the neural encoding of movements has been difficult to decode in the simple spikes of Purkinje cells (Helmchen and Buttner, 1995;Thier et al., 2000;Roitman et al., 2005;Roitman et al., 2009;Hewitt et al., 2011). As learning produces a change in the neural coding, this poor understanding has made it difficult to relate the rich behavioral changes that have been observed during motor learning, such as multiple timescales of memory (Smith et al., 2006;Kording et al., 2007), spontaneous recovery (Ethier et al., 2008), and savings (Pekny et al., 2011), with the neural mechanisms of learning in the cerebellum. Our long-term goal in building this marmoset lab is to search for the neural basis of learning internal models. However, the first step is to understand the neural code with which Purkinje cells make predictions.

Recently, using data in the macaque we proposed that the Purkinje cells may be organized in micro-clusters (Herzfeld et al., 2015;Herzfeld et al., 2018), wherein each micro-cluster is composed of Purkinje cells that share a common preference for error. The preference for error is expressed through the complex spike tuning of the Purkinje cell (Soetedjo and Fuchs, 2006;Soetedjo et al., 2008). We found that if Purkinje cells were organized in this way, the language of internal models as expressed by Purkinje cells became decipherable: simple spikes as a population predicted the real-time motion of the eye during a saccade (Herzfeld et al., 2015). Thus, this theory suggested a potential solution to the encoding problem: organize the simple spikes of a population of Purkinje cells based on the functional properties of each cell’s complex spikes.

If the neural representation of internal model relies on a population coding of Purkinje cells, the challenge is to simultaneously record simple and complex spikes from multiple Purkinje cells in the awake, behaving animals. However, this has proven to be difficult. Array electrodes have simultaneously isolated multiple Purkinje cells in rodents (Welsh et al., 1995;Sugihara et al., 2007;Blenkinsop and Lang, 2011;Tang et al., 2016;Tang et al., 2019) and decerebrated cats (Ebner and Bloedel, 1981), but this has often been in the anesthetized animal (however, see (Han et al., 2018)). In the awake behaving subject, calcium imaging has been used to track complex spikes in multiple Purkinje cells (Najafi et al., 2014;Kostadinov et al., 2019), but this technique does not have the temporal resolution yet to track simple spikes. In primates, simultaneous recordings from Purkinje cells are quite rare. Until our work, the report by Medina and Lisberger (2007) is the only example we are aware of in which pairs of Purkinje cells were recorded in the primate during goal-directed behavior.

### Limitations

Although the X-shaped head-post design had the advantage that the skin would cover nearly the entire skull, we found that within about a year following surgery the skin in subjects B and M gradually receded to the boundary of the dental cement. One of our challenges is to find ways to slow and eventually prevent this process of skin contraction, which might include experimentation with different forms of cement or its elimination.

Our experience with array type electrodes that have contacts along a single surface suggests that despite their high density, these electrodes can provide isolation of multiple Purkinje cells along neighboring folia, but rarely within a single folium. The goal of simultaneously recording from many Purkinje cells within a single folium will likely require employment of multiple electrodes.

Isolation of multiple Purkinje cells introduces the computational problem of spike attribution, as one contact can record complex spikes while the other records only simple spikes. Here, we showed that conditional probabilities that consider the effect of complex spikes on simple spikes can help with the attribution problem, but the task becomes much harder when many contacts are involved.

In summary, in order to better understand the neural basis of internal models, we have built a new laboratory that focuses on marmosets. Here, we reported on our attempts to improve marmoset behavioral training and electrophysiological recording methods. We found that through behavioral shaping, it was possible to motivate the animals to perform around 1000 trials in a head-fixed saccade task. We also found that with the aid of imaging, it was possible to build a system for simultaneous isolation of multiple Purkinje cells in the cerebellum.

## Acknowledgements

The work was supported by grants from the NIH (5-R01-NS078311), the Office of Naval Research (N00014-15-1-2312), and the National Science Foundation (CNS-1714623). We are grateful to Tahl Holtzman from Cambridge Neurotech, who made important suggestions regarding choice of microdrive, electrodes, and silicone gel for protection of the dura. We are also grateful to Mehrdad Jazayeri and Mark Churchland, who gave invaluable advice on design of the laboratory. ESN collected some of the behavioral data on subject B and all of the behavioral and neurophysiological data on subject M. He performed surgery on subjects M and R, analyzed the behavioral and neurophysiological data, and made figures. DJH designed and performed surgery, wrote the software for behavioral training, and collected behavioral data from subject B. PH designed the image guided procedures for construction of the head-post, designed the alignment ruler, and made figures. KK wrote some of the software, collected some of the behavioral data on subject M, and analyzed the neurophysiological data. TP performed image co-registration, made figures, and collected behavioral data on subject R. XW provided the animals as well as technical help with surgery and feedback for electrophysiology designs. RS collected some of the neurophysiological data and wrote the manuscript. We are very grateful to Seth Koehler, Xiao-Ping Liu, and Wang Lab technicians in helping with surgeries and animal care.

